# TPST-dependent and -independent regulation of root development and signaling by PSK LRR receptor kinases in Arabidopsis

**DOI:** 10.1101/2021.03.26.437140

**Authors:** Christine Kaufmann, Nils Stührwohldt, Margret Sauter

## Abstract

Tyrosine-sulfated peptides are key regulators of plant growth and development. The disulfated pentapeptide phytosulfokine (PSK) mediates growth via leucine-rich repeat receptor-like kinases, PSKR1 and PSKR2. PSKRs are part of a response module at the plasma membrane that mediates short-term growth responses, but downstream signaling of transcriptional regulation remains unexplored. In Arabidopsis, tyrosine sulfation is catalyzed by a single-copy gene (*TPST*). We performed a microarray-based transcriptome analysis in the *tpst-1* mutant background that lacks sulfated peptides to identify PSK-regulated genes and genes that are regulated by other sulfated peptides. Of the 160 PSK-regulated genes, several had functions in root growth and development in agreement with shorter roots and a higher lateral root density in *tpst-1*. Further, *tpst-1* roots developed higher numbers of root hairs and PSK induced expression of *WEREWOLF (WER)*, its paralog *MYB DOMAIN PROTEIN 23 (MYB23)* and *At1g66800* that maintain non-hair cell fate. The *tpst-1 pskr1-3 pskr2-1* mutant showed even shorter roots, and higher lateral root and root hair density than *tpst-1* revealing unexpected synergistic effects of ligand and PSK receptor deficiencies. While residual activities may exist, overexpression of *PSKR1* in the *tpst-1* background induced root growth suggesting that PSKR1 may be active in the absence of sulfated ligands.

**Highlight:** Phytosulfokine (PSK) receptor signaling promotes root elongation, determines lateral root density and maintains non-hair cell fate partially independent of TPST responsible for the activating sulfation of PSK.

## Introduction

Post-translationally modified peptides are an emerging class of signaling molecules that are involved in regulation of plant growth and development, and in responses to abiotic and biotic stresses including pathogen attacks. Post-translationally modified peptides are derived from larger pre-pro-proteins that require several steps of proteolytic cleavages. Besides proteolytic processing to release the peptide moiety from the inactive precursor, they may rely on tyrosine sulfation, proline hydroxylation and arabinosylation of hydroxyprolines to receive full peptide activities (Matsubayashi, 2014; Stührwohldt and Schaller, 2019; Kaufmann and Sauter, 2019).

Tyrosine sulfation of signaling peptides is catalyzed by the Golgi-localized tyrosylprotein sulfotransferase (TPST) that is encoded by a single-copy gene in Arabidopsis (Komori *et al.*, 2009). Five classes of sulfated peptides have so far been identified that include Phytosulfokine (PSK), Plant Peptides Containing Sulfated Tyrosine (PSYs), Root Meristem Growth Factors (RGFs, Golven (GLV) and CLE-like (CLEL)), Casparian Strip Integrity Factors (CIFs) and Twisted Seed 1 (TWS1). For most of them including PSK, experimental data indicated that tyrosine sulfation is required for full peptide activity (Matsubayashi and Sakagami, 1996; Matsubayashi *et al.*, 2006; Amano *et al.*, 2007; Kutschmar *et al.*, 2009; Matsuzaki *et al.*, 2010; Whitford *et al.*, 2012; Meng *et al.*, 2012; Fernandez *et al.*, 2013; Doblas *et al.*, 2017; Nakayama *et al.*, 2017).

TPST loss-of-function mutants are an important tool to analyze functions of and processes triggered by tyrosine-sulfated peptides. Several mutants have been identified all lacking activity of TPST and consequently activities of tyrosine-sulfated peptides (Komori *et al.*, 2009; Zhou *et al.*, 2010; Kang *et al.*, 2014). *tpst* mutants, known as *tpst-1*, *active quiescent center* (*aqc1-1* to *aqc1-3*) and *hypersensitive to Pi starvation7* (*hps7*), are characterized by an overall dwarfed phenotype, pale-green leaves and stunted roots which are brought about by defective maintenance of the root stem cell niche, decreased meristematic activity and decreased cell expansion (Komori *et al.*, 2009; Zhou *et al.*, 2010; Kang *et al.*, 2014) and hypersensitivity to fructose (Zhong et al., 2020). TPST acts to maintain the root stem cell niche by regulating basal and auxin-induced expression of the transcription factors Plethora 1 and 2 (PLT1 and PLT2) (Zhou *et al.*, 2010).

Phytosulfokine is one of the most extensively studied sulfated peptides with regards to physiological functions. Mature PSK is a disulfated pentapeptide of the sequence (Y(SO_3_H)-I-Y(SO_3_H)-T-Q) that is derived from precursor proteins that vary in length from 77 to 109 amino acids in Arabidopsis (Kaufmann and Sauter, 2019). Contrary to all other sulfated peptides, PSK precursor proteins share a fully conserved sequence of the mature pentapeptide with PSK6, that differs in the last amino acid of the PSK pentapeptide, as an exception. However, since no expressed sequence tags have been reported and expression levels according to RNA-seq data are extremely low, *PSK6* is considered to be a pseudogene (Kaufmann and Sauter, 2017).

The enzymes that are responsible for precursor processing to release the PSK pentapeptide are largely unknown. The subtilisin-like serine protease 1.1 (SBT1.1) was shown to cleave the Arabidopsis PSK4 precursor peptide *in vitro* (Srivastava *et al.*, 2008), however the cleavage site is N-terminal of the mature peptide within the variable part of the precursor, and the physiological relevance of this cleavage and of SBT1.1 for PSK biogenesis remains unclear. In tomato, an aspartate-specific SBT (tomato phytaspase 2, *S*lPhyt2) was shown to be involved in PSK maturation (Reichardt *et al.*, 2020). The eight PSK precursors in tomato (Reichardt *et al.*, 2020), and the seven PSK precursors in Arabidopsis (Kaufmann and Sauter, 2019) share an aspartate residue on the amino side of the PSK pentapeptide. However, the Arabidopsis protease cleaving the N-terminal Asp has not been identified to date. An unusual precursor protein is PSK1 that is flanked by Asp at both sites of the mature peptide. We could recently show that the C-terminal Asp is cleaved by Arabidopsis SBT3.8 to release the active C-terminus (Stührwohldt *et al.*, 2021).

PSK was originally identified as a growth factor that promotes cell division of Asparagus cells grown in culture at low density (Matsubayashi and Sakagami, 1996) and has since been linked to multiple physiological functions. In plant reproductive processes, PSK promotes pollen germination (Chen *et al.*, 2000), pollen tube elongation (Stührwohldt *et al.*, 2015) and it guides the pollen tube from the transmitting tract along the funiculus to the embryo sac to support seed production (Stührwohldt *et al.*, 2015). Further, PSK induces root, hypocotyl and leaf growth mainly by promoting cell expansion (Kutschmar *et al.*, 2009; Stührwohldt *et al.*, 2011; Hartmann *et al.*, 2014). Cotton fiber elongation is also driven by enhanced cell elongation and promoted by overexpression of a putative *GhPSK* gene (Han *et al.*, 2014). In addition, PSK differentially affects plant immunity and stress responses. It supports the response to the necrotrophic fungi *Alternaria brassicicola* and *Sclerotinia sclerotiorum* as well as the bacterium *Ralstonia solanacearum* and it represses the response to hemi-/biotrophs such as *Pseudomonas syringae* and the oomycete *Hyaloperonospora arabidopsidis* (Loivamäki *et al.*, 2010; Igarashi *et al.*, 2012; Mosher *et al.*, 2013; Rodiuc *et al.*, 2016). Zhang et al. (2018) showed that PSK signals the Auxin-dependent immune responses in tomato after infection with the necrotrophic fungus *Botrytis cinerea*. Recent studies also revealed a role of PSK signaling in osmotic and drought stress adaptation (Schönhof *et al.*, 2020; Stührwohldt *et al.*, 2021).

The major understanding of PSK signaling comes from the identification of the plasma membrane-localized PSKR receptors PSKR1 and PSKR2, that belong to the large family of leucine-rich repeat receptor-like kinases (LRR-RLKs). PSK binds extracellularly to the island domain of PSKR1 and PSKR2 located between the leucine-rich repeats (Matsubayashi *et al.*, 2006; Wang *et al.*, 2015). The intracellular PSKR1 domain functions as an essential kinase (Irving *et al.*, 2012; Hartmann *et al.*, 2014; Hartmann *et al.*, 2015; Kaufmann and Sauter, 2017). At defined Ser, Thr and Tyr residues, the PSKR1 kinase autophosphorylates (Hartmann *et al.*, 2015; Mitra *et al.*, 2015; Muleya *et al.*, 2016; Kaufmann and Sauter, 2017). PSKR1 forms a heterodimer together with the promiscuous co-receptor BAK1/SERK3, a SERK family member (Ladwig *et al.*, 2015; Wang *et al.*, 2015), directly interacts with the H^+^-ATPases AHA1 and AHA2, and forms a functional complex with Cyclic Nucleotide Gated Channel 17 (CNGC17) (Ladwig *et al.*, 2015).

While the response module consisting of PSK receptors, BAK1, AHA1, AHA2 and CNGC17 induces PSK-mediated growth, only a few downstream components of PSK signaling are known (Zhang et al., 2018). To identify new players in PSK signaling, we set up a microarray approach to identify PSK-regulated genes and genes that are regulated by other sulfated peptides by using the *tpst-1* mutant as a tool. The transcriptome data prompted a more detailed analysis of root development. The results revealed a role of PSKR signaling in determining lateral root density, and in maintaining non-hair cell fate by regulating the transcription factor genes *WEREWOLF (WER)*, its paralog *MYB DOMAIN PROTEIN 23* (*MYB23*) and *At1g66800* (Lee and Schiefelbein, 1999; Matsui *et al.*, 2005; Deal and Henikoff, 2010). The characterization of a *tpst-1 pskr1-3 pskr2-1* triple mutant revealed unexpected synergistic effects of TPST deficiency and PSK receptor deficiency that may suggest activity of PSK receptors dependent and independent from its ligand PSK.

## Results

### Identification of genes differentially regulated by PSK

TPST is encoded by a single-copy gene in Arabidopsis. In *tpst* knock-outs, production of sulfated peptides is abolished leading to a sulfated ligand-free background. The *tpst-1* loss-of-function mutant of Arabidopsis has been a useful tool to study control of root development by sulfated peptide hormones (Komori *et al.*, 2009). Root growth induction by enhanced cell expansion is one of the best-characterized functions of the disulfated pentapeptide PSK (Matsubayashi *et al.*, 2006; Kutschmar *et al.*, 2009). In order to identify genes that are regulated by PSK and to compare these to genes that are regulated by other sulfated peptides, we performed a microarray experiment on roots of five-day-old wild-type, *tpst-1* and *tpst-1* seedlings that were treated with 1 μM PSK (Figure 1A). In total, we identified 615 genes that were differentially regulated between wild-type and the *tpst-1* mutant (FC≥2; Figure 1B). When we compared roots of *tpst-1* seedlings that were supplemented with PSK to roots of untreated *tpst-1* seedlings, 240 genes were found to be differentially regulated. The comparison between *tpst-1* seedlings treated with PSK and wild-type seedlings resulted in 265 differentially expressed genes (Figure 1B). The overlap of genes differentially regulated by PSK in *tpst-1* seedling roots and of genes differentially regulated between *tpst-1* and wild type was determined to identify genes which are specifically regulated by PSK and not by other sulfated peptides (Figure 2A). This comparison revealed 169 genes that are specifically regulated by PSK.

**Figure 1:**
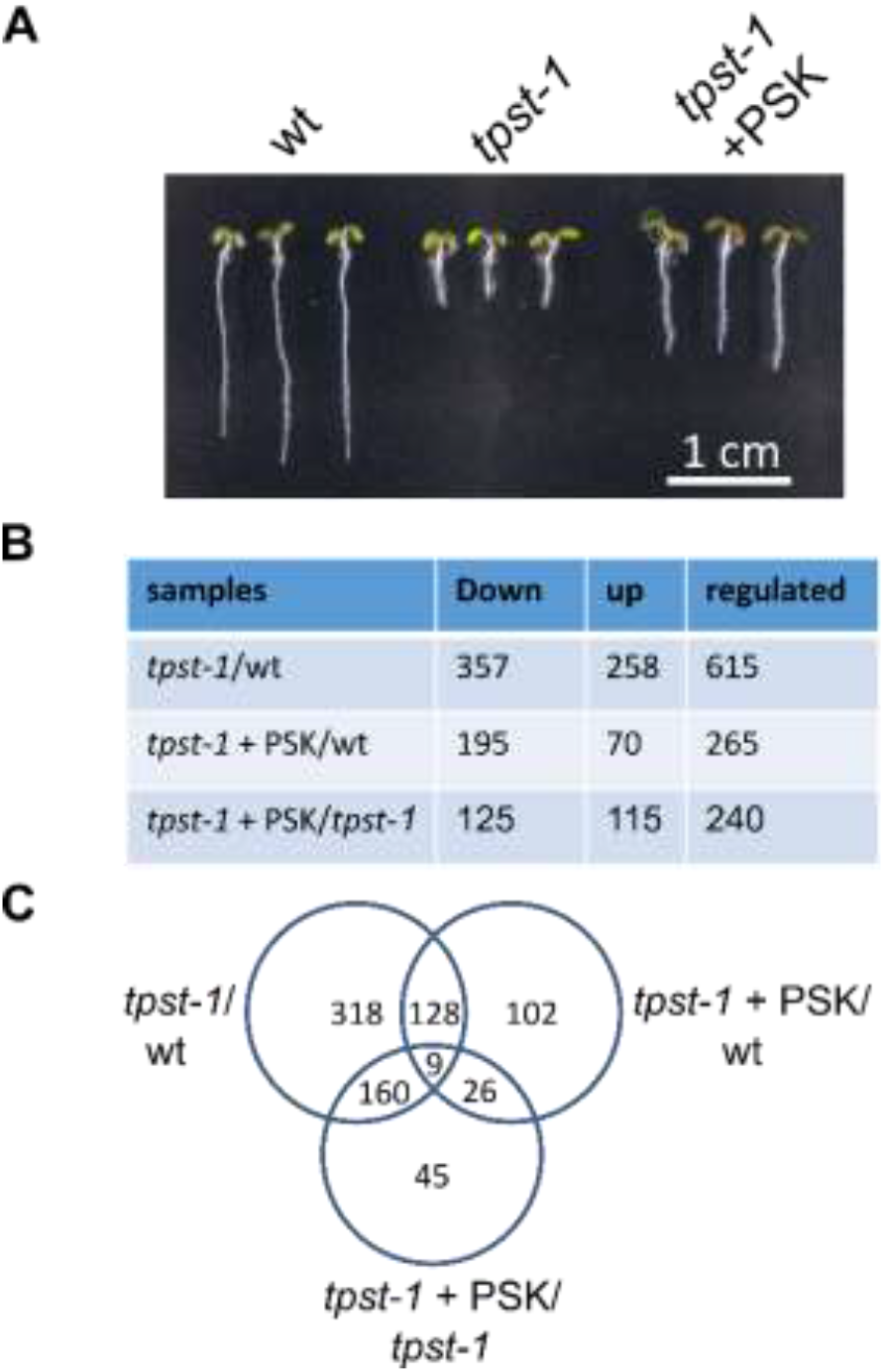
Transcriptomic analysis of genes regulated by sulfated peptides. **(A)** Representative seedlings of wild type, *tpst-1* and *tpst-1* treated with 1 μM PSK grown for five days under long-day conditions. Complete roots were harvested and material was subjected to microarray analysis. Scale bar represents 1 cm. **(B)** Table with numbers of down- and upregulated genes and the total number of genes regulated. The different samples compared are *tpst-1* versus wild type, *tpst-1* + PSK versus wild type and *tpst-1* + PSK versus *tpst-1*. Microarray experiment was performed with three biological replicates each. A cutoff at a *P* value of 0.001 was used to indicate differentially expressed genes combined with a cutoff at a fold change of two. **(C)** Venn diagram of genes regulated between the different genotypes and treatments. Total numbers of genes regulated are given.

**Figure 2:**
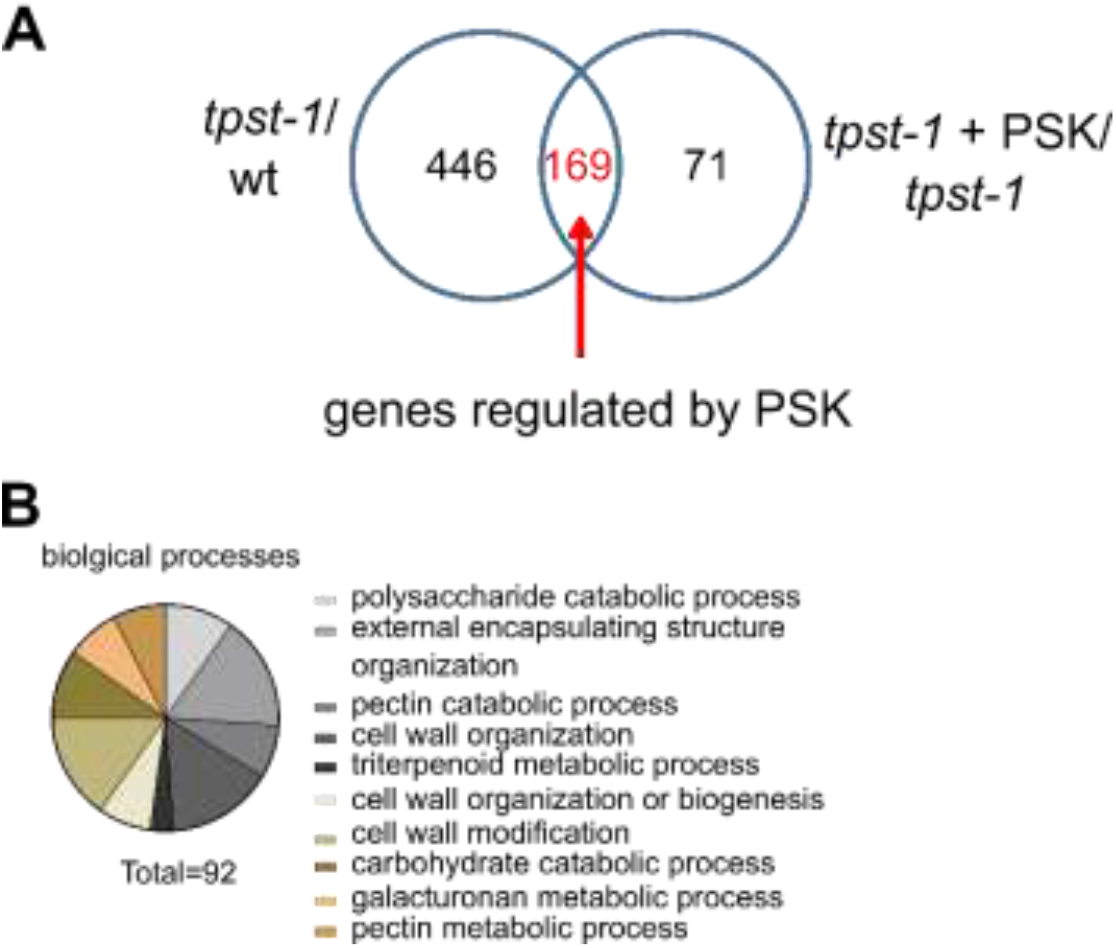
Identification of (overrepresented) genes regulated by PSK. **(A)** Venn diagram that illustrates the number of genes regulated in the *tpst-1* versus wild type and *tpst-1* + PSK versus *tpst-1*. The overlap between these samples identifies the number of genes that are regulated by PSK. **(B)** Biological processes that are overrepresented among the genes that are regulated by PSK. The identified genes were analyzed by the PANTHER16.0 program (Mi *et al.*, 2019; Mi *et al.*, 2021). A total of 92 genes could be assigned to specific, overrepresented biological processes.

PSK regulates hypocotyl and root growth mainly by promoting cell expansion (Kutschmar *et al.*, 2009; Stührwohldt *et al.*, 2011). Cell expansion requires cell wall remodeling which includes changes in cell wall composition and structure. To categorize the genes that are regulated by PSK we analyzed the biological processes that are overrepresented with the publicly available database PANTHER (Mi *et al.*, 2019; Mi *et al.*, 2021). Here, we observed that PSK regulates genes overrepresented in biological processes related to cell wall modification, organization or biogenesis which includes pectin and galacturonan metabolism as well as carbohydrate catabolism (Figure 2B). Our data link gene expression changes induced by PSK signaling to growth triggered by PSK (Figure 2A, B). We also detected an overrepresentation of genes linked to triterpenoid metabolism (Figure 2B). Activity of one of the genes, *Marneral Synthase 1* (*MRN1*), has previously been linked to cell elongation (Go *et al.*, 2012).

To independently verify differential gene expression in response to PSK, we selected six candidate genes and analyzed their relative transcript levels by RT-qPCR (Figure 3). We tested three genes each that were down- or upregulated by PSK (Figure 3A). For *MRN1* (*At5g42600*), *Terpene Synthase-Like 23* (*TPS23*, *At3g25820*) and *Thalian-diol Desaturase* (*THAD1*, *At5g47990*) which encode for terpene biosynthesis enzymes (Figure 3A), we confirmed positive regulation by PSK (Figure 3 B-D). Of three genes that were negatively regulated by PSK according to the microarray experiment, a receptor-like kinase (*At5g41290*), a basic helix-loop-helix transcription factor (*BHLH129*, *At2g43140*) and *Baruol Synthase 1* (*Bars1*, *At4g15370*), we confirmed downregulation by PSK for the *receptor-like kinase* and *BARS1* (Figure 3E-G). To test for short- and long-term regulation of genes by PSK, we exposed *tpst-1* seedlings in hydroponic culture to 100 nM PSK for 4, 8, 12, 16, 20, 24 and 48 hours or kept them as controls without PSK (Supplemental Figure S1). Genes of baruol, marneral and thalianol synthesis are organized in clusters and genes within a cluster were coordinately regulated (Field and Osbourn, 2008). For the three baruol biosynthesis genes *BARS1*, *CYP705A3* and *CYP705A2* identified in the microarray, we observed significant downregulation of transcripts over time with the highest reduction in expression after 48 hours (Supplemental Figure S1). Expression of the three marneral biosynthesis genes and of the three thalianol genes increased over time in response to PSK suggesting that the microarray data reliably revealed genes differentially regulated by PSK (Supplemental Figure S1).

**Figure 3:**
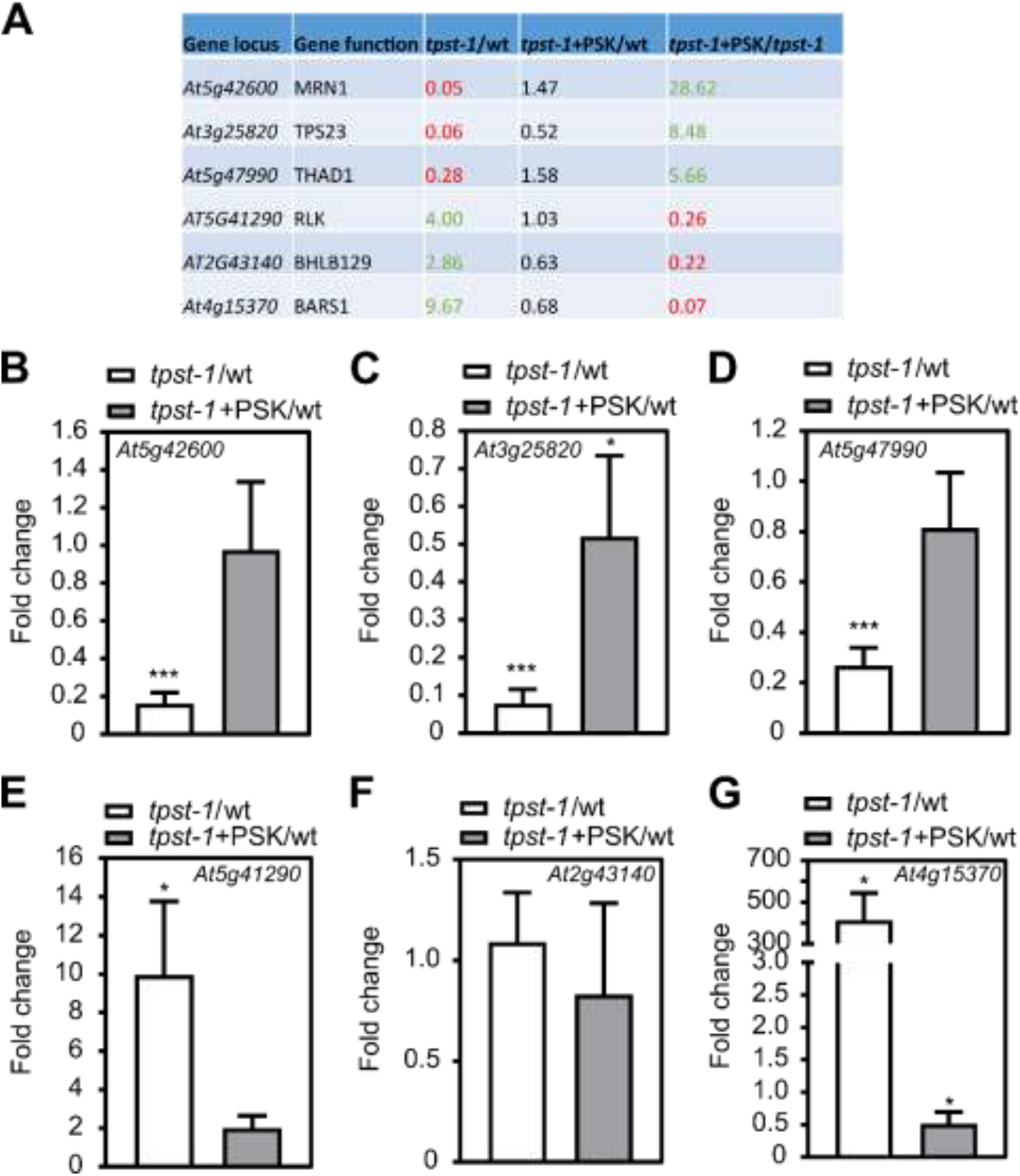
Verification of PSK-regulated genes in Arabidopsis roots by qPCR. **(A)** Table of chosen genes identified as PSK-regulated within the microarray analysis. Six genes were chosen that were regulated by PSK, but not significantly regulated in *tpst-1* + PSK versus wild type. **(B-G)** Fold change of expression tested by qPCR of **(B)** *At5g42600 (MRN1)* **(C)** *At3g25820 (TPS23)* **(D)** *At5g47990 (THAD1)* **(E)** *At5g41290* (a receptor-like kinase) **(F)** *At2g43140 (BHLH129)* and **(G)** *At4g15370 (BARS1)*. Roots were harvested from seedlings grown for 5 days under control conditions or treated with 1 μM of PSK. qPCR was performed on three biological replicates with two technical repeats, and gene expression was normalized to two reference genes. Results are shown as fold changes in *tpst-1* versus wild type and *tpst-1* + PSK versus *tpst-1*. Each time point included pooled plant material of several independent seedlings. * and *** indicate significant differences compared to the control at P < 0.05 or P < 0.001, respectively (two-tailed t test).

### The *tpst-1 pskr1-3 pskr2-1* triple mutant shows an enhanced phenotype compared with *tpst-1*

Genes that were regulated similarly in *tpst-1* versus wild type and *tpst-1* treated with PSK versus wild type were defined as regulated by sulfated peptides other than PSK. Here, we identified 128 genes (Supplemental Figure 2A). Putative candidate peptides responsible for the regulation of these genes are PSYs, RGFs, CIFs and TWS1. The biological processes that were overrepresented in this group included responses to reactive oxygen species, reactive nitrogen species, salicylic acid and metal and iron ions (Supplemental Figure S2B) in accord with recently published data that showed control of root meristem size by RGF1 through ROS signaling (Yamada *et al.*, 2020). Overall, genes overrepresented in the *tpst-1* mutant could be assigned to cell wall modifications, reactive oxygen species, nitric oxide and secondary metabolism (Supplemental Figure S3).

Tyrosine sulfation is a prerequisite for activity of sulfated peptides. Consequently, loss of peptide signaling in the *tpst-1* mutant is complemented by the addition of sulfated peptides (Komori *et al.*, 2009; Matsuzaki *et al.*, 2010; Doblas *et al.*, 2017). The PSK receptor double knockout mutant *pskr1-3 pskr2-1*, is insensitive to PSK for instance with regard to root growth promotion (Kutschmar *et al.*, 2009) (Figure 4A, B). The analysis of the gene expression data revealed regulation of genes that we could not clearly categorize as being regulated by PSK or other sulfated peptides (see examples in Supplemental Figure S4). Aside from genes that were regulated in *tpst-1*, but were not regulated by PSK (category A) we identified genes that were regulated in *tpst-1* with only partial restoration of expression by PSK (category B) and genes that were only differentially expressed when comparing *tpst-1* supplemented with PSK to wild type (category C) (Supplemental Figure S4) suggestive of crosstalk between TPST-dependent signaling pathways. To be able to more clearly address PSK-dependent root developmental processes in the sulfated peptide-deficient background, we created a triple mutant by crossing the *tpst-1* mutant with the PSK receptor double mutant, *pskr1-3 pskr2-1*. Knockout of alle three genes was confirmed by RT-PCR (Supplemental Figure S5A). We expected that the *tpst-1 pskr1-3 pskr2-1* triple mutant should phenocopy the *tpst-1* mutant but should be insensitive to PSK as is the *pskr1-3 pskr2-1* mutant. Unexpectedly, the combined knockout of PSK receptors and *TPST* genes had synergistic effects (Figure 4A, B). The *tpst-1 pskr1-3 pskr2-1* and *pskr1-3 pskr2-1* seedling, were insensitive to PSK as expected (Figure 4A, B). But whereas the primary roots of *pskr1-3 pskr2-1* and *tpst-1* seedlings were reduced in length by 25.5% and 73.5%, respectively, *tpst-1 pskr1-3 pskr2-1* triple mutant roots were reduced by 87.4% which is significantly more than in *tpst-1* (Figure 4A, B). Synergistic effects in the *tpst-1 pskr1-3 pskr2-1* triple mutant compared to *tpst-1* and *pskr1-3 pskr2-1* were also observed with regard to overall plant architecture and rosette size (Figure 4C, D).

**Figure 4:**
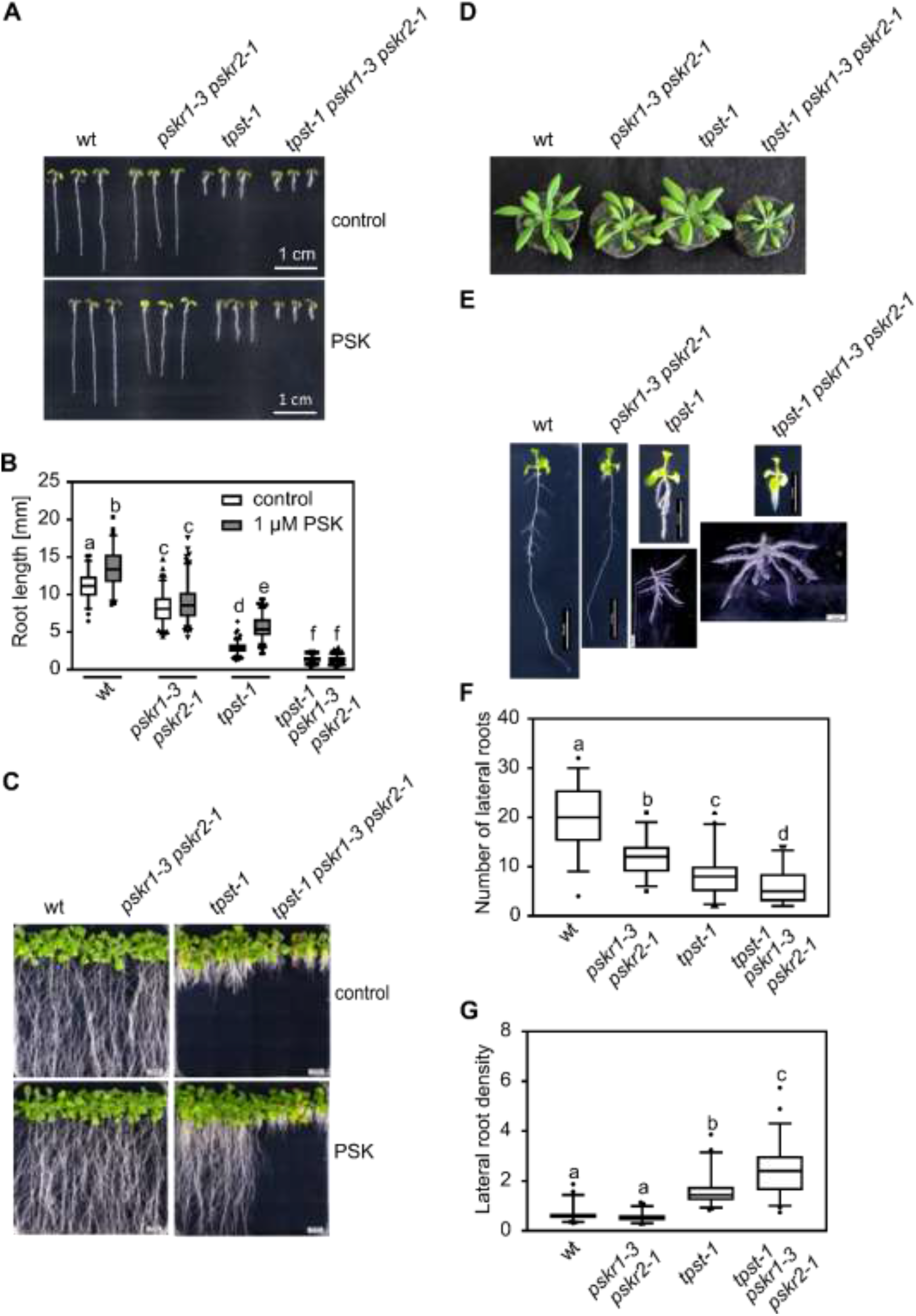
A *tpst-1 pskr1-3 pskr2-1* triple mutant shows an unexpected, synergistic phenotype. **(A)** Representative seedlings of wild type, *pskr1-3 pskr2-1*, *tpst-*1 and *tpst-1 pskr1-3 pskr2-1* grown with or without 1 μM of PSK for five days under long-day conditions. Scale bar indicates 1 cm. **(B)** Root length [mm] of 5-day-old wild type, *pskr1-3 pskr2-1*, *tpst-*1 and *tpst-1 pskr1-3 pskr2-1* treated without or with 1 μM of PSK. **(C, D)** Representative images of respective genotypes grown **(C)** for three weeks under sterile conditions with and without PSK or **(D)** for four weeks on soil under long-day conditions. **(E)** Representative images of plants grown for 11 days under sterile growth conditions. For *tpst-1* and *tpst-*1 and *tpst-1 pskr1-3 pskr2-1* a representative example of lateral root development is shown. Scale bars represent the indicated lengths. **(F)** Number of lateral roods and **(G)** lateral root density of 11-day old plants of wild type, *pskr1-3 pskr2-1*, *tpst-*1 and *tpst-1 pskr1-3 pskr2-1*. The root and the shoot were separated and lateral roots were spread to determine initiation sites of lateral roots. Experiments were performed at least three times with similar results. Data are shown for one representative experiment as the mean ± SE, n ≥ 68, (F) n ≥ 46, (G) n ≥ 46. Different letters indicate significant differences (Kruskal-Wallis, *P*<0.05).

The observation that several genes related to lateral root development were regulated in the *tpst-1* mutant (Supplemental Figure S6), prompted us to analyze lateral roots in the *tpst-1*, *pskr1-3 pskr2-1* and the *tpst-1 pskr1-3 pskr2-1* mutants. Lateral root density in *pskr1-3 pskr2-1* seedlings was comparable to that in wild type whereas the number of lateral roots was significantly reduced likely due to a shorter primary root (Figure 4 E-G) (Kutschmar *et al.*, 2009). The *tpst-1* mutant had a significantly reduced number of lateral roots compared to wild type and *pskr1-3 pskr2-1* whereas lateral root density was increased reporting that sulfated peptides other than PSK control lateral root density. In *tpst-1 pskr1-3 pskr2-1* seedlings these phenotypes were even more pronounced with fewer lateral roots and a higher lateral root density than in *tpst-1* seedlings (Figure 4F, G). These findings raised the hypothesis that lateral root development could, to some extent, be triggered by PSK receptor activity independent of the sulfated ligand PSK. Likewise, it is conceivable that residual receptor activity exists in *pskr1-3 pskr2-1* and signaling via these residual receptor activity is saturated by endogenous PSK in accord with the observation that the *pskr1-3 pskr2-1* mutant is insensitive to exogenous PSK (Kutschmar et al., 2009). Also, residual peptide sulfation activity cannot be completely excluded in *tpst-1*. However, *tpst-1* shows a 30-fold higher sensitivity toward exogenous PSK compared to wild type indicating that PSK receptors are largely in an unbound state (Stührwohldt et al., 2011). In this case, expressing additional receptors should not promote a PSK response.

To address this issue, we overexpressed *PSKR1* or *PSKR2* in the *tpst-1* mutant background (Figure 5, Supplemental Figure S5B, C) under the control of the constitutive 35S promoter. Overexpression of *PSKR1* or *PSKR2* was verified by semi-quantitative RT-PCR for two and three lines, respectively (Supplemental Figure S5B, C) and the lines were analyzed for primary root elongation as an easy-to-monitor readout. Strikingly, overexpression of *PSKR1* in the *tpst-1* background induced root growth (Figure 5A, B) while *PSKR2* overexpression did not (Figure 5C, D) suggesting that PSKR1 has growth-promoting activity in the absence of sulfated ligands. Root elongation of the PSKR1 overexpressors in the *tpst-1* background was comparable to that of *tpst-1* seedlings treated with PSK and was not further promoted by addition of PSK (Figure 5A, B). These findings indicated that lack of sulfated PSK can be compensated for by increasing the abundance of PSKR1; lines with high PSKR1 abundance have saturated PSKR1 signaling independent from its ligand (Figure 5A, B).

**Figure 5:**
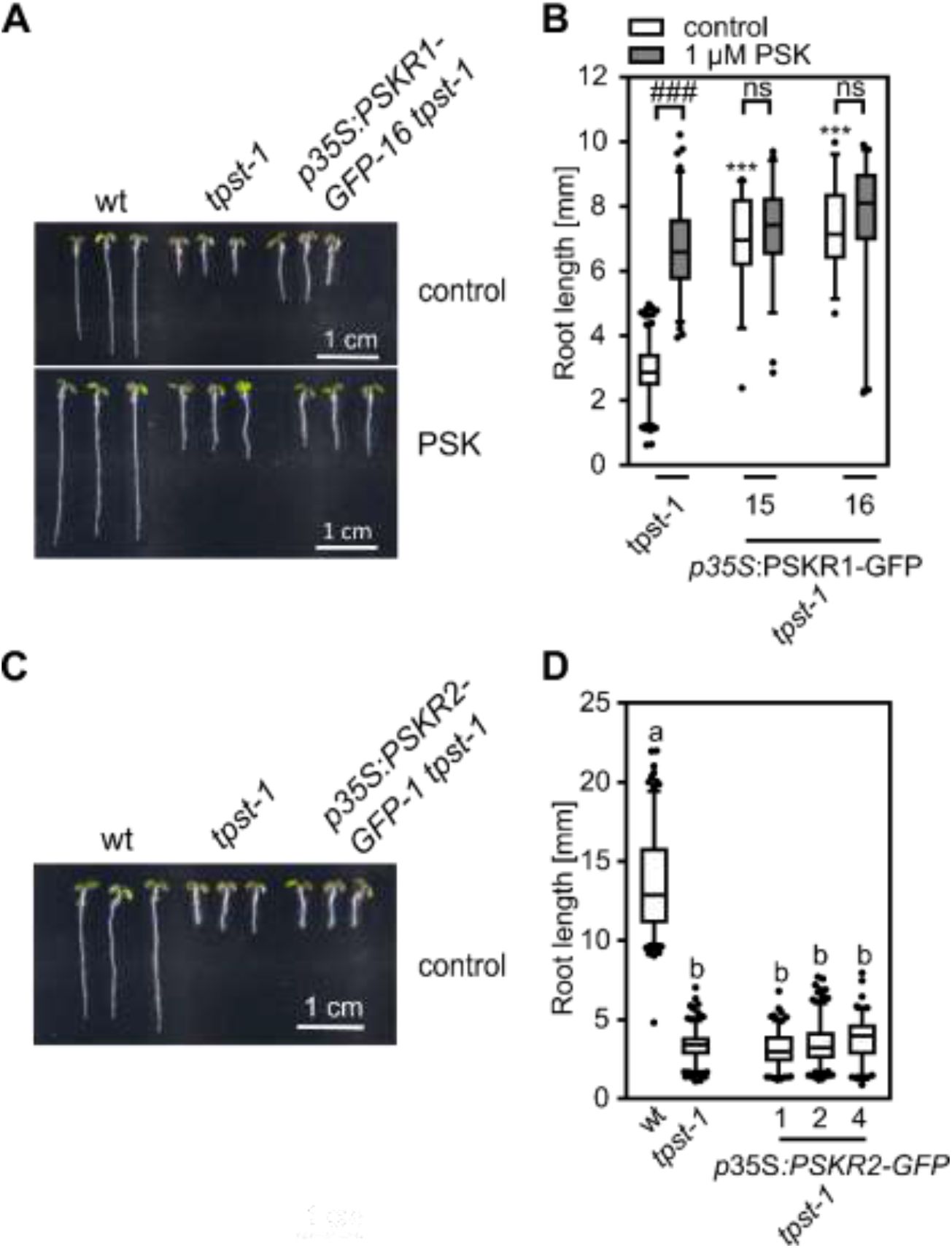
Overexpression of *PSKR1* in the *tpst-1* background promotes root growth. **(A, C)** Representative seedlings of wild type, *tpst-1*, *p35S*:PSKR1-GFP *tpst-1* and *p35S*:PSKR2-GFP *tpst-1* grown for 5 days without (A. C) and with 1 μM PSK (A). **(B, D)** Root lengths [mm] of **(B)** wt, *tpst-1* and two independent lines of *p35S*:PSKR1-GFP *tpst-1* and **(D)** wt, *tpst-1* and *p35S*:PSKR1-GFP *tpst-1* supplemented without (B, D) or with 1 μM PSK (B). Numbers indicate independent transgenic lines. (B) Experiments were performed at least three times with similar results. Data are shown for one representative experiment as the mean ± SE. (D) Experiment was performed once. Data are shown as the mean ± SE. (B) n ≥ 31, (D) n ≥ 122. In (B), *** indicates significant differences to the wild-type control at P < 0.001 (two-tailed t test). ^###^ indicates a significant difference in comparison to the untreated control at P < 0.001 (two-tailed t test). In (D), different letters indicate significant differences (Kruskal-Wallis, *P*<0.05). In (A) and (C), scale bars represent 1 cm.

### PSKRs signal non-hair cell fate through WER expression

We found that the *hypersensitive to Pi starvation 7* (*hps7)* mutant, that is allelic to *tpst-1*, shows a root hair phenotype that has however not been addressed previously (Kang *et al.*, 2014). Furthermore, genes involved in the control of root hair formation are differentially regulated by PSK based on our microarray data (Figure 6A). We therefore asked whether TPST and PSK receptors play a similar synergistic role in root hair formation as they do in root growth and lateral root formation. *tpst-1* displays an abnormal root hair phenotype (Figure 6B). Root hair formation in the *tpst-1 pskr1-3 pskr2-1* triple mutant was even more pronounced (Figure 6B). To analyze root hair repression by the TPST-PSKR signal pathway in more detail, we transformed wild-type, *tpst-1*, *pskr1-3 pskr2-1,* and *tpst-1 pskr1-3 pskr2-1* plants with the trichoblast-specific reporter *pEXPANSIN7*:*β*-glucuronidase (*pEXP7:GUS*) (Cho and Cosgrove, 2002) to visualize and quantify root hairs (Figure 6C). Promoter activity excluding the meristematic zones was detected in roots of all genotypes (Figure 6C).

**Figure 6:**
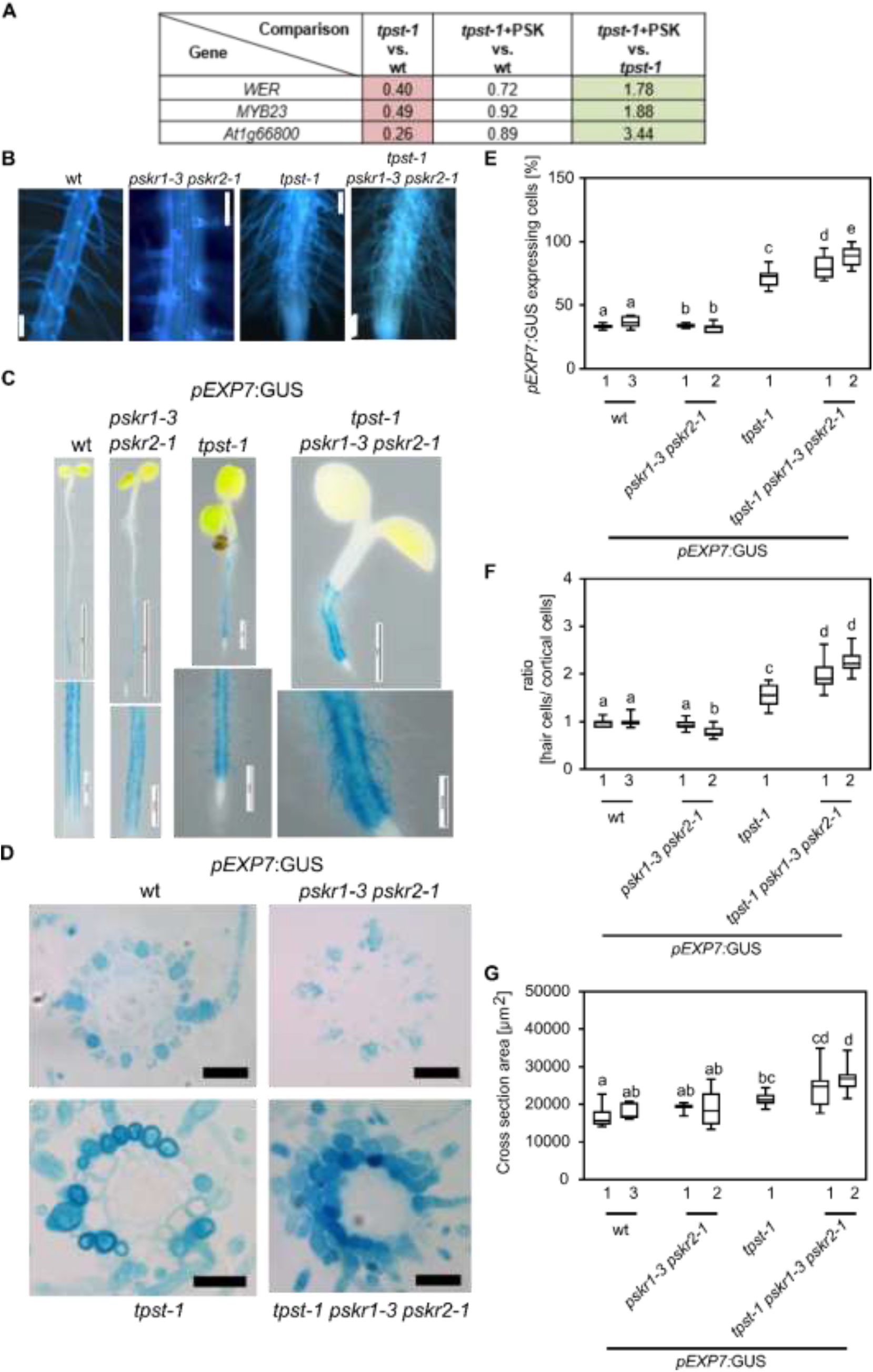
TPST and PSKRs control root hair formation by altering position-dependent epidermal cell fate determination. **(A)** Table with fold change of *WER*, *MYB23* and *At1g66800* identified from the microarray experiment. Microarray experiments were performed with three biological replicates. **(B)** Auto-fluorescence imaging of the root hair zone of wild type, *pskr1-3 pskr2-1, tpst-1* and *tpst-1 pskr1-3 pskr2-1*. The scale bar represents 100 μm. **(C)** Representative images of five-day-old wild-type, *pskr1-3 pskr2-1, tpst-1* and *tpst-1 pskr1-3 pskr2-1* seedlings expressing *pEXP7*:*GUS*. Scale bars represent the indicated lengths. **(D)** Representative cross sections of 5-day-old *pEXP7:GUS* seedlings; scale bar represents 50 μm. **(E)** Quantification of cells expressing *pEXP7:GUS* as a marker for trichoblasts. **(F)** Ratio of *pEXP7:GUS*-expressing hair cells to cortical cells in genotypes indicated. **(G)** Cross section area [μm^2^] determined from cross sections of *pEXP7p*:*GUS*-expressing seedlings. Numbers indicate independent transgenic lines. Experiments were performed at least three times with similar results. Data are shown for one representative experiment as the mean ± SE, (E) n ≥ 9, (F) n ≥ 9, (G) n ≥ 9. Different letters indicate significant differences (Kruskal-Wallis, *P*<0.05).

To determine the percentage of *pEXP7:GUS* expressing cells, the ratio of hair cells to cortical cells and the cross section areas of the mutant roots, we made cross sections from the root hair zone (Figure 6D). In two *pEXP7:GUS pskr1-3 pskr2-1* lines, the percentage of cells with root hair identity was 34.1 and 32.4%, respectively, compared to 33.5% and 36.8% in two *pEXP7:GUS* wild-type lines (Figure 6E). In *pEXP7:GUS tpst-1* seedlings, 71.9% of epidermal cells had root hair cell identity indicating that sulfated peptide signaling determines non-hair cell fate (Figure 6E). In *pEXP7:GUS tpst-1 pskr1-3 pskr2-1* seedlings, the number of root hair cells was significantly higher than in *pEXP7:GUS tpst-1* seedlings with 79.5% and 88.4% of epidermal cells expressing the hair cell marker (Figure 6E) indicating that root hair formation was suppressed by sulfated peptide signaling, in part via PSK receptors.

Hair cell formation in Arabidopsis is determined by the position of epidermal cells with regard to the cortical cell layer (Ma *et al.*, 2001; Salazar-Henao *et al.*, 2016). Hair cells touch two cortex cells while non-hair cells border on a single cortex cell (Figure 7A). To gain more insight into the activity of TPST and PSKRs in the position-dependent control of root hair formation, we analyzed hair and cortical cells in more detail. While the root cross section area increased significantly in *tpst-1* and *tpst-1 pskr1-3 pskr2-1* compared to wild type (Figure 6G), the number of epidermal cells remained constant in all four genotypes (Supplemental Figure S7A). Further, the number of cortex cells only increased slightly in the mutants (Supplemental Figure S7B). Consequently, we determined the ratio of hair cells to cortex cells that was 1:1 in wild type and in *pskr1-3 pskr2-1* (Figure 6F). The ratio of hair to cortex cells increased to 1.5:1 in *tpst-1* and to 2.1:1 in *tpst-1 pskr1-3 pskr2-1*. The increase in root hair cells in *tpst-1* and *tpst-1 pskr1-3 pskr2-1* was not a result of more hair (H)-positions above two cortical cells, but rather due to loss of position-dependent determination of epidermal cell fate with trichoblasts developing at non-hair (N)-positions as illustrated in Figure 7A and experimentally shown for *tpst-1* and *tpst-1 pskr1-3 pskr2-1* in Figure 7B.

**Figure 7:**
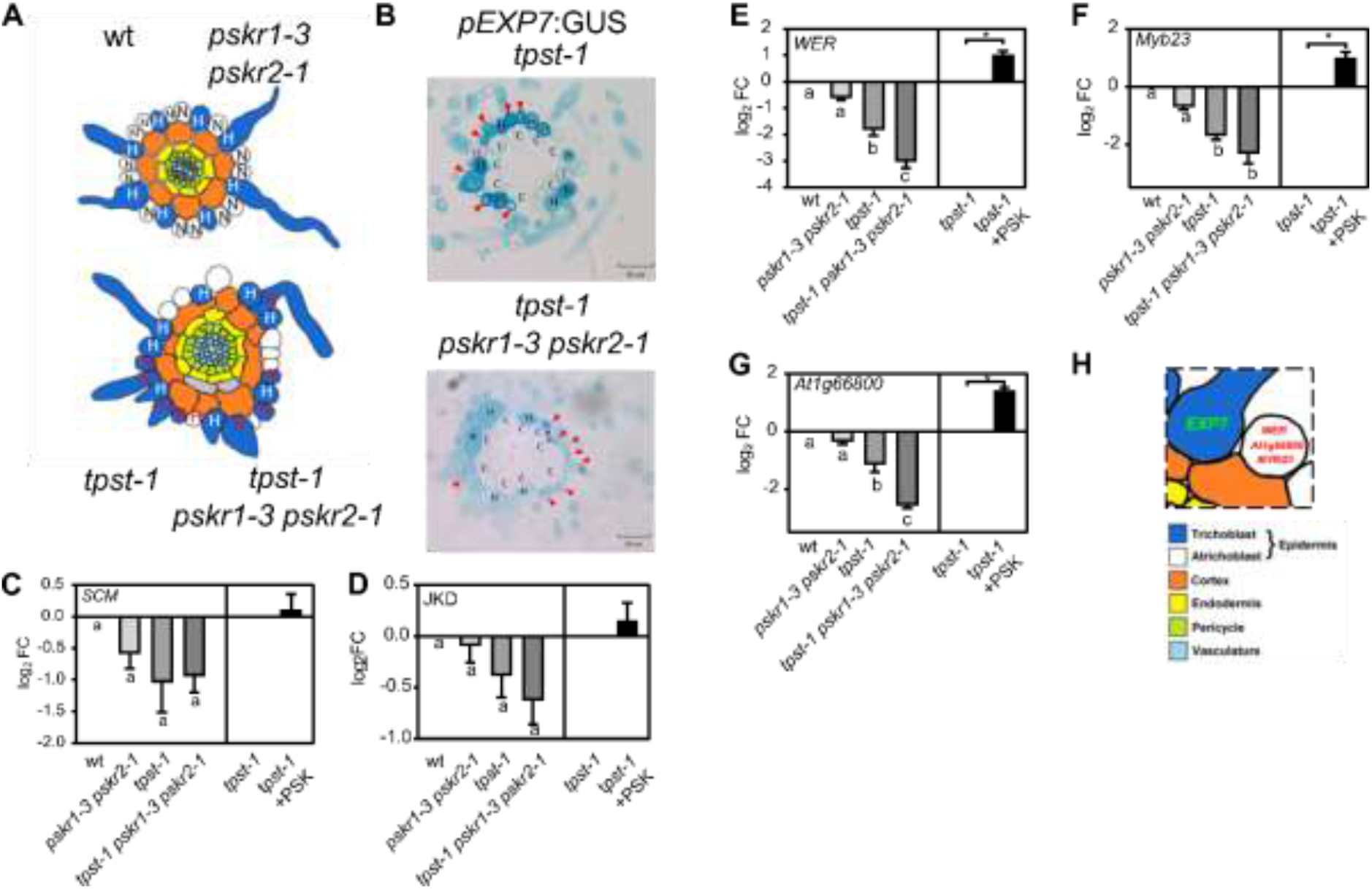
TPST and PSKRs control root hair formation by regulation of three marker genes for non-hair cell fate. **(A)** Schematic presentation indicating the position of trichoblasts and cortical cells. Trichoblasts are in an H position in wild type and *pskr1-3 pskr2-1*. In *tpst-1* and *tpst-1 pskr1-3 pskr2-1*, they are developed in H and N positions. **(B)** Representative cross sections of 5-day-old *tpst-1* and *tpst-1 pskr1-3 pskr2-1* seedlings expressing *pEXP7:GUS*. Arrows indicate *pEXP7:GUS*-expressing cells in the N (non-hair root) position. Red H marks a cell that is in an H (root hair) position, but does not express *pEXP7:GUS.* Scale bars indicate 50 μm **(C, D, E, F, G)** Log_2_ fold change of expression tested by qPCR of non-hair cell fate marker genes **(C)** SCM (LRR receptor-like kinase Scrambled), **(D)** JKD (Jackdaw), **(E)** WER (Werwolf) **(F)** Myb23 and **(G)** At1g66800. Seedlings were grown for 5 days under control conditions or treated with 1 μM of PSK and subjected to RT-qPCR analysis. RT-qPCR was performed on three biological replicates with two technical repeats, and gene expression was normalized to two reference genes and is shown as log_2_ fold change. Each time point included pooled plant material of several independent seedlings. Asterisks indicate significant differences between *tpst-1* and *tpst-1* treated with PSK. Different letters indicate significant differences (Kruskal-Wallis, *P*<0.05). **(H)** Schematic presentation indicating expression of marker genes *EXP7*, *WER*, *MYB23* and *At1g66800* in Arabidopsis wild-type seedlings.

To identify the downstream targets of the TPST/PSKR1 signals that determine non-hair cell fate, we tested expression of two major regulators of non-hair cell fate, the transcription factor JACKDAW (JKD) (Hassan *et al.*, 2010), expressed in cortical cells, and the LRR-RLK SCRAMBLED (SCM) localized at the plasma membrane of trichoblasts. Expression of either gene was not significantly different between wild-type, *pskr1-3 pskr2-1*, *tpst-1* or *tpst-1 pskr1-3 pskr2-1* roots (Figure 7C, D). Further, PSK did not induce the expression of *JKD* or *SCM* indicating that they are not transcriptional targets of TPST/PSKR signaling (Figure 7C, D). Next, we tested expression of the downstream atrichoblast-specific transcription factors WER, MYB23 and At1g66800. In our microarray experiment, all three transcription factors were downregulated in the *tpst-1* mutant compared to the wild type, and induced by PSK, for *WER* and *MYB23* by a fold change of 1.66 and 1.78, respectively (Figure 6A). Expression analysis by RT-qPCR showed that all three transcription factors were slightly, but not significantly reduced in the *pskr1-3 pskr2-1* mutant (Figure 7E-G). However, all three genes were significantly downregulated in *tpst-1* versus wild type and further repressed in *tpst-1 pskr1-3 pskr2-1* compared to *tpst-1* (Figure 7E-G) again revealing a synergistic effect of the mutations. Expression of *WER*, *MYB23* and *At1g66800* was induced by PSK in *tpst-1* indicating that PSKR signaling maintains non-hair cell fate with PSK enhancing the signal output (Figure 7E-G). In conclusion, PSK acts downstream or independent of JKD and SCM and promotes expression of *WER*, *MYB23* and *At1g66800*. In this pathway, PSKR may act dependent and independent of its ligand PSK to promote non-hair cell fate (Figure 7H).

## Discussion

### Transcriptional crosstalk between peptide signaling pathways

The use of the *tpst-1* mutant for our transcriptome analysis turned out as a helpful approach to identify genes that are regulated by PSK in roots. A detailed expression analysis of triterpenoid synthesis genes showed that differential gene regulation detected in steady-state expression profiles by microarray analysis in *tpst-1* seedlings grown in the presence or absence of PSK, in many cases successfully predicted gene regulation also in the short term as shown in time course analysis by RT-qPCR. Growth of above-ground parts of plants overexpressing thalianol synthase (THAS) or MRN1 was significantly inhibited indicating that an excess amount of thalianol, marnerol, or their derivatives might be detrimental for shoot growth whereas plants overexpressing THAS have longer roots than the wild type or *thas* knock-out lines suggesting a root-specific growth promoting effect (Field and Osbourn, 2008; Field *et al.*, 2011).

The *tpst-1* mutant, when compared to wild type, revealed genes regulated by sulfated peptides in general. For some genes, regulation by several sulfated peptides seems likely. Some of the genes that were identified as regulated in *tpst-1* were only partially regulated by PSK indicating crosstalk between different sulfated peptide signaling pathways (Zhou et al., 2010, Whitford et al., 2012, Zhang et al., 2018). Common gene targets and crosstalk between different signaling peptide pathways agrees with the observation that root growth, that is stunted in *tpst-1*, is fully restored by the joint application of PSK, PSY1 and RGF1 (Matsuzaki *et al.*, 2010). Joint application of PSK and PSY1 restores cell-elongation activity, but does not affect meristematic activity in *tpst-1*, as evidenced by meristem size (Matsuzaki *et al.*, 2010). Single peptides partially rescue the short-root phenotype whereas PSK and RGF1 together promote root growth in a non-additive manner (Matsuzaki *et al.*, 2010), indicating crosstalk between the signaling pathways. The transcriptome approach taken here provides a repertoire of data and allows the assignment of genes that are regulated by PSK, other sulfated peptides, or genes regulated by PSK and other sulfated peptides.

### Downstream targets of PSK and PSK receptor signaling in root growth

*TPST* is highly expressed in lateral root primordia (Komori *et al.*, 2009), is induced by auxin and affects the expression of *PLT1/2*, *PIN* genes, and auxin biosynthetic genes (Zhou *et al.*, 2010). Since lateral roots are induced by auxin (Péret *et al.*, 2009) these findings agree with a role of TPST in the control of lateral root density. Overexpression of several Golven (RGF, CLEL) peptides in Arabidopsis reduced lateral root density (Fernandez *et al.*, 2013) indicating that Golven peptides regulate lateral root architecture jointly with other sulfated peptides, e.g. PSK or PSY1, possibly at different developmental stages.

In addition to primary root elongation and lateral root formation, root system architecture is regulated by Tyr-sulfated peptides through the control of root hair development. Sulfated peptides Golven 4 (RGF7, CLEL4) and Golven 8 (CLEL5) have been implicated in root hair elongation as indicated by shorter root hairs in lines where peptides were silenced or knocked out (Fernandez *et al.*, 2013). In *tpst-1*, more root hairs develop indicating that sulfated signaling peptides act to maintain nonhair cell fate. Aside from the *tpst-1* T-DNA insertion line (Komori *et al.*, 2009), other *TPST* mutants have been identified, the active quiescent center mutants (*aqc1-1* to *aqc1-3*) (Zhou *et al.*, 2010) and the *hypersensitive to Pi starvation 7* (*hps7)* mutant (Kang *et al.*, 2014). The *hps7* mutant displayed a similar root hair phenotype as *tpst-1* that was exaggerated by phosphate starvation with root hair formation extending toward the tip close to the meristem revealing a link between TPST-dependent signaling in nutrient stress adaptation. Support for a broader contribution of sulfated peptide signaling in abiotic stress responses comes from recent reports on the role for PSK in mediating osmotic stress tolerance (Stührwohldt *et al.*, 2021).

In root hair development, TPST-dependent signals appear to act downstream of the transcription factor JKD and the kinase SCM. Both participate in the cortex-to-epidermis cell communication, and ensure that root hairs form from epidermal cells that are in contact with two cortex cells but not from epidermal cells that are in touch with a single cortex cell leading to natural spacing of root hair cell files in Arabidopsis (Hassan *et al.*, 2010). JKD and SCM act upstream of the transcription factors WER and MYB23 that are expressed atrichoblast-specifically (Schiefelbein *et al.*, 2014). SCM represses *WER* transcripts, preferentially in hair cells. WER and MYB23 are key repressors of root hair identity in epidermal cells that, when knocked out, lead to a hairy root phenotype (Grierson et al., 2014; Matsui et al., 2005). Strikingly, atrichoblast-specific expression of PSKR1 is sufficient to promote root growth (Hartmann *et al.*, 2013) providing an unexpected spatial link between PSK-receptor activity, promotion of root elongation and inhibition of root hair formation. The involvement of PSKR1 signaling in root hair is recorded by the reinforced effects of physiological and transcriptional effects in the *tpst-1 pskr1-3 pskr2-1* triple mutant.

While expression of *JKD* and *SCM* was not altered in *tpst-1*, transcript levels of *WER* and *MYB23* were reduced compared to wild type and were at least partially restored by PSK suggestive of a role of sulfated peptide signaling in suppression of trichoblast differentiation. In *tpst-1* seedlings, the control of hair cell identity by cortex cells is lost and root hairs develop at non-hair positions. Together these findings indicate that protein sulfation by TPST is a crucial element in cell-to-cell communication that helps to establish cell identities in the root epidermis.

### Ligand-(in)dependent regulation of receptor activities

Hormone receptors are considered to be activated through binding of their respective ligand (Hohmann *et al.*, 2017). High-affinity binding of PSK to the ectodomain of its receptor depends on Tyr-sulfation of the PSK pro-protein by TPST (Wang *et al.*, 2015). PSK-binding to PSKR1 at the island domain within the ectodomain is predicted to promote binding to the co-receptor BAK1/SERK3 or other members of the SERK family (Ladwig *et al.*, 2015; Hohmann *et al.*, 2017), but experimental evidence is lacking that the PSKR/BAK1 heterodimerization is dependent on PSK binding. Mutual phosphorylations between receptor and co-receptor activate the cytosolic PSKR kinase and initiate signaling. The soluble kinase domain of PSKR1 displays auto- and transphosphorylation activities (Hartmann *et al.*, 2015; Kaufmann and Sauter, 2017). Among other residues, two Ser residues in the juxtamembrane region are autophosphorylated by the soluble PSKR1 kinase. This phosphorylation not only occurs *in vitro*, but was demonstrated to also occur *in planta* on the full-length receptor (Kaufmann and Sauter, 2017). Site-directed mutagenesis of the phosphorylated Ser residues altered substrate phosphorylation activity *in vitro* and shoot growth *in planta* indicating that the soluble kinase domain of PSKR1 acquires an active conformation. Root growth promotion by overexpression of PSKR1 in the PSK-deficient *tpst-1* background suggests that the receptor can similarly acquire an active state in the absence of a sulfated ligand either by an unsulfated ligand, receptor modification or interaction with other proteins.

ABA signaling in plants is crucial for plant adaptation to dehydration which became important when plants conquered terrestrial habitats. ABA signaling relies on Pyrobacter resistant/Pyrobacter resistant-like (PYR/PYL) ABA receptor-induced repression of PP2C phosphatases which leads to the activation of downstream SNF-related kinases. While most algae do not possess an ABA receptor, it was recently reported that an algal PYR/PYL-homolog encoded by *Zygnema circumcarinatum*, a member of the algae sister lineage of land plants, represses PP2C activity independent of ABA suggesting that the signaling function of the ABA receptor evolved prior to ligand binding (Sun *et al.*, 2019). In case that a receptor and a ligand did not strictly co-evolve, it is conceivable that residual ligand-independent receptor signaling is retained. It is conceivable that ligand and PSK receptors evolved independently early in land plant evolution and that residual ligand-independent signaling was retained (Furumizu *et al.*, 2020). A recently characterized temperature-sensitive mutation in the FERONIA (FER) receptor that recognizes rapid alkalinization factor (RALF), is point-mutated in a conserved Gly outside the ligand binding site in ectodomain of the receptor (Kim *et al.*, 2021). An analogous mutation in the related RALF receptor THESEUS1 (THE1) leads to loss of receptor function while retaining ligand binding (Hématy *et al.*, 2007; Gonneau *et al.*, 2018). The findings confirm that receptor activity is not exclusively dependent on ligand binding. Whether the mutations in FER and THE1 interfere with a conformational change required for receptor activation remains speculative.

PSKR1 and related LRR-RLKs such as BRI1 interact in multimeric protein complexes, the composition of which may differ depending on cell type, physiological state and environmental signals. No studies have yet been done to evaluate the impact of the protein composition of the PSK receptor module on receptor output but it is conceivable that the phosphorylation status and protein interactions impact the conformation and activity of the intracellular kinase Analysis of the *tpst-1 pskr1-3 pskr2-1* mutant revealed a strong synergistic effect of ligand and receptor knockout in all three root phenotypes analyzed, primary root elongation, lateral root formation and root hair formation. These observations seem hardly explained by residual receptor and ligand activities. More so, overexpression of *PSKR1* in the ligand-free *tpst-1* background increased root length arguing for a ligand-independent mode of receptor activation or for basal ligand-independent receptor activity. These observations should stimulate research focused on the ligand-dependent and ligand-independent composition of protein complexes and their signal output in different cell types and under changing environmental conditions. Such high-resolution analyses might help to shed light on the unexplored paths of ligand-independent receptor signaling.

## Materials and Methods

### Growth conditions and plant material

All experiments were carried out with *Arabidopsis thaliana* ecotype Col-0. The T-DNA insertion lines *tpst-1* (SALK_009847) and the double knock out line *pskr1-3 pskr2-1* were described previously (Komori *et al.*, 2009; Kutschmar *et al.*, 2009; Stührwohldt *et al.*, 2011). The triple knockout line *tpst-1 pskr1-3 pskr2-1* was generated by crossing *tpst-1* with *pskr1-3 pskr2-1*. Loss of all three transcripts was verified by semi-quantitative RT-PCR. Seeds were surface-sterilized in 2% (v/v) sodium hypochlorite for 15 min, washed five times with autoclaved water and subsequently laid out on 0.5 x Murashige-Skoog medium (Duchefa, Harlem, Netherlands), 1.5% (w/v) sucrose, solidified with 0.4% (w/v) Gelrite (Duchefa Harlem, Netherlands). If indicated, media were supplemented with 1 μM PSK (Pepscan, Lelystad, Netherlands). For growth on soil, plants were grown in a 2:3 sand:humus mixture that was frozen at −80°C for 2 days to avoid insect contamination and watered regularly with tap water. After two days of stratification at 4°C in darkness, plants were transferred to long-day conditions (16 h light with 70 μM photons m^−2^ s^−1^, 8 h dark) at 21-22°C and 60% humidity for the times indicated.

### Cloning of constructs and generation of transgenic lines

To generate a reporter for trichoblasts, the previously described 437 base pair-long promoter region from −386 to +48 of *EXPANSIN7* (*At1g12560*) (Cho and Cosgrove, 2002) was amplified using the forward primer 5’-ACGCGCGGCCGCGTGTTCAATTTAACTAATCATTG-3’ with a cleavage site for *Not*I and the reverse primer 5’-ACGCCTCGAGCTATTGAGAAGAATTTAAAGCT-3’ with an *Xho*I cleavage site and ligated into pENTR1a DS to generate the *pEXP7:GUS* reporter. The construct was sequenced and recombined into pBGWFS7 by using the Gateway cloning system (Thermo Fisher Scientific, Waltham, Massachusetts, USA). Cloning of the *p35S:PSKR1-GFP* construct into pB7WG2.0 has been described previously (Hartmann *et al.*, 2013). Plant transformation and selection of transgenic plants was done as described (Kaufmann *et al.*, 2017).

### Preparation and analysis of cross sections and GUS staining

*pEXP7:GUS* expressing seedlings of the genetic backgrounds indicated were grown on plates for 5 days, collected and stained with GUS staining solution (Vielle-Calzada *et al.*, 2000). Roots were separated from shoots and embedded in TechnoVit (Heraeus Kulzer, Wehrheim, Germany) as described in the manufacturer’s manual. Cross sections of 10 μm thickness were prepared with a Leica RM 2255 microtome, collected on glass slides, embedded in CV Mount solution (Leica, Bensheim, Germany) and analyzed with an Olympus BX41 microscope. Pictures were taken with an Infinity 3S camera using the software Infinity Analyze 6.5 (Lumenera, Ottawa, Canada). The numbers of epidermal and cortical cells were counted on pictures of cross sections. The cross section area was determined with Fiji/ImageJ open-source software (https://imagej.net/Fiji) from the same pictures.

### RNA isolation and gene expression analyses

For microarray and RT-qPCR analyses, roots from five-day-old seedlings that were grown on plates supplemented with or without 1 μM PSK under sterile condition were used. Total RNA was isolated with TRI-reagent (Sigma Aldrich, St. Louis, USA) following manufacturer’s instructions. RNA was dissolved in DEPC-treated H_2_O and the quality and quantity of RNA was measured with a NanoDrop spectrometer (ThermoFisher Scientific, Waltham, Massachusetts, USA). For RT-PCR and RT-qPCR, 1 μg mRNA was digested with DNaseI and subsequently reverse-transcribed with OligodT primers. Quantitative PCR was performed with the Rotor-Gene SYBR Green PCR Kit (Qiagen, Venlo, Netherlands) according to manufacturer’s instructions. The reverse transcription products were amplified using gene-specific primers as indicated in Supplementary Table 1. Reactions were performed with a Rotor Gene Q cycler (Qiagen, Venlo, Netherlands). Data (takeoff and efficiencies) was given by ‘Comparative quantification analysis’ from the cycler-corresponding Rotor Gene Q Series software (Qiagen, Venlo, Netherlands). The fold change was calculated by normalization to the geometric mean of *ACT2* and *GAPC* expression. For statistical analysis log2-transformed fold change values were used. At least three independent biological replicates with technical repeats each were performed.

Microarray experiments were performed with three biological replicates of three samples each, wt, *tpst-1* and *tpst-1* treated with 1 μM PSK. AraGene-1_0-st; Affymetrix microarray slides containing 38408 transcripts were used for transcriptome analysis. Analysis of RNA quality, chip hybridization, and data processing were performed at the MicroArray Facility (Flanders Institute for Biotechnology (VIB), Leuven, Belgium). Briefly, analysis was based on the RMA expression values. To identify differentially expressed genes, the Robust Multi-Array Average (RMA) expression values at the different conditions were compared with the LiMMA package of Bioconductor (Wettenhall and Smyth, 2004; Smyth, 2005). For each contrast of interest, it was tested if it deviated significantly from 0 with a moderated *t* statistic implemented in LiMMA. The resulting *P* values were corrected for multiple testing with Benjamini-Hochberg to control the false discovery rate (Benjamini and Hochberg, 1995). All *P* values given were corrected for multiple testing. A cutoff at a *P* value of 0.001 was used to indicate differentially expressed genes combined with a cutoff at a fold change of two.

### Statistical analysis

Data sets were analyzed for normal distribution. In case of normal distribution for all-pairwise comparison, an ANOVA with α=0.05 was run with Origin 8.5 software, whereas data sets that were not normally distributed were analyzed with a Kruskal– Wallis test with Bonferroni as *P*-value adjustment method (α=0.05) by using the package “agricolae” [65] and statistics software R (https://CRAN.R-project.org/doc/FAQ/R-FAQ.html).

## Author contributions

CK, NS and MS designed experiments. NS performed initial characterization of *tpst-1 pskr1-3 pskr2-1* and performed microarray experiment. CK performed all other experiments and analyses. NS and MS wrote the manuscript with contributions from CK.

## Acknowledgement

Authors thank Timo Staffel (Plant Developmental Biology and Physiology, University of Kiel) for excellent technical support.

## Supplemental Figure Legends

**Supplemental Figure 1: Expression analysis of genes involved in triterpene synthesis in response to PSK treatment.** log_2_ fold chance of expression of **(A)** baruol, **(B)** marneral and **(C)** thalianol biosynthesis genes. Expression was tested in roots of *tpst-1* seedlings that were grown for five days in hydroponic culture and treated with or without 100 nM PSK for the time indicated. Small or capital letters indicate significant differences between time points of control samples or PSK-treated samples (ANOVA, post-hoc Bonferroni, p<0.05), whereas asterisks indicate significant difference between treatments at a specific time point (Student t-test; p<0.05; n=3 three).

**Supplemental Figure 2: Genes regulated by other sulfated peptides than PSK. (A)** Venn diagram of genes regulated between the different genotypes and treatments indicated. Total number of genes regulated are given. The red number indicates the number of genes regulated by other sulfated peptides than PSK **(B)** Biological processes that are overrepresented among the genes that are regulated by other sulfated peptides than PSK. The identified genes were analyzed by the PANTHER16.0 program (Mi *et al.*, 2019; Mi *et al.*, 2021). A total of 42 genes could be assigned to biological processes that are overrepresented.

**Supplemental Figure 3: Regulated genes overrepresented in the** *tpst-1* **mutant.** Biological processes that are overrepresented among the genes that are regulated in the *tpst-1* versus wild type. The identified genes were analyzed by the PANTHER16.0 program (Mi *et al.*, 2019; Mi *et al.*, 2021). A total of 102 genes could be assigned to biological processes that are overrepresented.

**Supplemental Figure 4: Genes that could not be clearly categorized as regulated by PSK or other sulfated peptides and were sorted by different categories.** Genes are derived from the microarray experiment. Microarray experiments were performed with three biological replicates.

**Supplemental Figure 5: Semi-quantitative RT-PCR analysis** of **(A)** wt and *tpst-1 pskr1-3 pskr2-1*, **(B)** wt, *tpst-1* and two independent *p35S*:PSKR1-GFP *tpst-1* lines and (C) wt, *tpst-1* and three independent *p35S*:PSKR2-GFP *tpst-1* lines. Transcript levels of *TPST*, *PSKR1*, *PSKR1-GFP* and/ or *PSKR2* and *PSKR2-GFP* were analyzed in 5-day-old seedlings by RT- PCR. *Actin2* was amplified as a control for RNA input.

**Supplemental Figure 6: Differentially regulated genes with functions in lateral root growth and development.** Genes are derived from the microarray experiment. Microarray experiments were performed with three biological replicates.

**Supplemental Figure 7: Analysis of epidermal and cortical cell numbers (A, B)** Quantification of (A) epidermal cell numbers and (B) cortical cell numbers in wild type, *pskr1-3 pskr2-1, tpst-1* and *tpst-1 pskr1-3 pskr2-1* that were determined from cross sections of *pEXP7:GUS*-expressing expressing lines. Numbers indicate independent transgenic lines. Experiments were performed at least three times with similar results. Data are shown for one representative experiment as the mean ± SE, (A) n ≥ 9, (B) n ≥ 9. Different letters indicate significant differences (Kruskal-Wallis, *P*<0.05).

## Supplemental Tables

**Supplemental Table 1: Primers used in this study**.

**Supplemental Table 2: Differentially expressed genes identified by microarray analysis**.

